# Connectome-based brain fingerprints predict early cognitive decline in Parkinson’s patients with minor hallucinations

**DOI:** 10.1101/2025.04.17.649310

**Authors:** Sara Stampacchia, Fosco Bernasconi, Dimitri Van De Ville, Enrico Amico, Javier Pagonabarraga, Jaime Kulisevsky, Olaf Blanke

## Abstract

Individual variability in connectome organization offers a unique framework for capturing patient-specific alterations and advancing personalized models in medicine. Minor hallucinations (MH) affect up to 40% of Parkinson’s disease (PD) patients and are early indicators of cognitive decline and dementia, hence crucial for early intervention. While previous studies focused on group-level differences, connectome-based brain fingerprinting enables deeper, individualized analysis of neural change. Applying this approach to PD patients with and without MH using resting-state fMRI, we show that each patient exhibited unique brain fingerprint, revealing rich quantifiable personalized features with medical relevance. MH-patients showed a loss of subject-specific features in brain networks linked to cognitive health, while somatosensory regions – typically less distinctive – became more prominent, emphasizing their role in MH pathogenesis. These differences enabled to identify – in an entirely data driven manner – patient-specific networks linked to early subclinical cognitive alterations, as well differential spatial fingerprinting organization linked to cortical densities of neurotransmitters. These findings reveal a distinct, patient-specific connectomic signature that differentiates PD patients with MH, uncovering early neural markers for precision medicine in PD.

## Introduction

Parkinson’s Disease (PD) is an irreversible, progressive neurodegenerative disease characterized by Lewy bodies (misfolded α-synuclein) ^1^, affecting the brainstem, nigrostriatal dopaminergic neurons, and other subcortical and cortical structures ^2^. While primarily a movement disorder, PD is characterized by early non-motor symptoms, including hallucinations ^3,4^, which are highly prevalent ^5–8^, affecting up to 50% of patients regularly ^9–11^, and with a prevalence of up to 70% in the advanced stages ^12,13^. Hallucinations are linked to faster cognitive decline, dementia ^9,14–17^, earlier home placement ^18–20^, and increased mortality risk ^18,20^.

Most studies on the neural substrates of hallucinations in PD have focused on complex visual hallucinations, the most common type in advanced PD ^14^. Activation studies suggest altered bottom-up and top-down visual processing in visual hallucinations, with reduced activation in primary visual cortices and increased activation in fronto-parietal regions during visual stimulation ^21^ and hallucinatory episodes ^22^. Similarly, resting-state functional connectivity studies have reported impaired coupling between attentional and visual networks, which may contribute to hallucinatory episodes ^23^. Moreover, the attentional network hypothesis ^24–26^ suggests that hallucinations in PD arise from decreased activation of the dorsal attentional network (DAN) when processing ambiguous visual stimuli arising from altered bottom-up processing in early visual areas ^27^, leading to over-reliance on top-down signals from default mode (DMN) and/or salience (SAL) networks.

Although the majority of past imaging work has focused on visual hallucinations in PD, so-called ‘minor’ hallucinations (MH) – including presence hallucinations, passage hallucinations, and pareidolias ^28,29^ –are of clinical relevance, because they often occur at earlier stages of PD ^11,30^, may even precede parkinsonian motor symptoms ^31^ and occur in up to 40% of patients ^9,32^. Recent work implicated the temporo-parietal junction and inferior frontal gyrus in robotically-induced MH ^33^, increased frontal EEG activity ^9^, and gray matter reduction in visuo-temporal regions in PD patients, further associated with mild cognitive impairment. Brain alterations observed in patients with MH partially overlap with those identified in visual hallucinations, including increased connectivity within the DMN ^34,35^, reduced DAN connectivity, and disrupted interactions between the DAN and the DMN^34^. Nevertheless, this body of research has primarily relied on group-averaged analyses, which do not capture the individual-specific nature of functional brain architecture—also known as the brain fingerprint ^36,37^. Considering the heterogeneity and individually distinct phenomenological expression of hallucinations, accounting for neural variability is particularly crucial.

Functional connectivity patterns estimated from resting-state functional magnetic resonance imaging (fMRI) or magnetoencephalography data – referred to as functional connectomes (FCs) – are unique to each individual ^36–38^. These *brain-fingerprints* have been linked to cognitive traits ^39,40^ and are altered in psychiatric ^41–43^ as well as neurodegenerative diseases, including Alzheimer’s Disease ^44,45^ and PD ^46,47^. While electrophysiological studies have reported preserved connectome identifiability in relation to motor impairment in PD ^46,47^, no study to date has explored this phenomenon using fMRI— leveraging its higher spatial resolution—or investigated its role in differentiating PD patients with and without hallucinations.

If functional connectomes are unique to each individual, they may also encode features relevant for personalized approaches, offering insights into clinically meaningful variability. Given the heterogeneous nature of hallucinations and their potential link to cognitive decline, it remains unclear whether functional brain fingerprints differ across early-stage PD patients with distinct clinical trajectories, namely PD with minor hallucinations (PD-MH) and PD without hallucinations (PD-nH). In this study, we investigate whether connectome-based fingerprints can identify patient-specific functional networks linked to clinical traits, with the goal of detecting, at an early stage, networks in PD-MH associated with cognitive impairment. To address this, we examined: (1) the preservation of fMRI brain fingerprints in PD, (2) their ability to distinguish between PD-MH and PD-nH, (3) their association with clinical traits and (4) their relationship with neurotransmitters systems implicated in hallucinations.

## Results

Our approach (see Methods) involved three steps: (1) We estimated functional connectomes (FCs) for each subject in the first and the second half of the fMRI acquisition ^45^(Fig. 1A). (2) We then assessed for each half brain fingerprints using identifiability matrices (Amico & Goñi, 2018a), deriving metrics such as *ISelf* (self-similarity), *IOthers* (similarity to others), *IDiff* (brain discriminability), and Success-rate ^36^ (Fig. 1B). (3) We analysed spatial specificity by computing individual FC-edge distinctiveness using intraclass correlation (ICC), mapping brain fingerprints across functional networks and onto the cortical surface (Fig. 2). (4) We finally tested whether functional connections with high fingerprinting capability would predict clinical scores (Fig. 3) and (5) explored the association between cortical topography of brain-fingerprints and neurotransmitters systems implicated in hallucinations (Fig.4).

**Figure 1.**
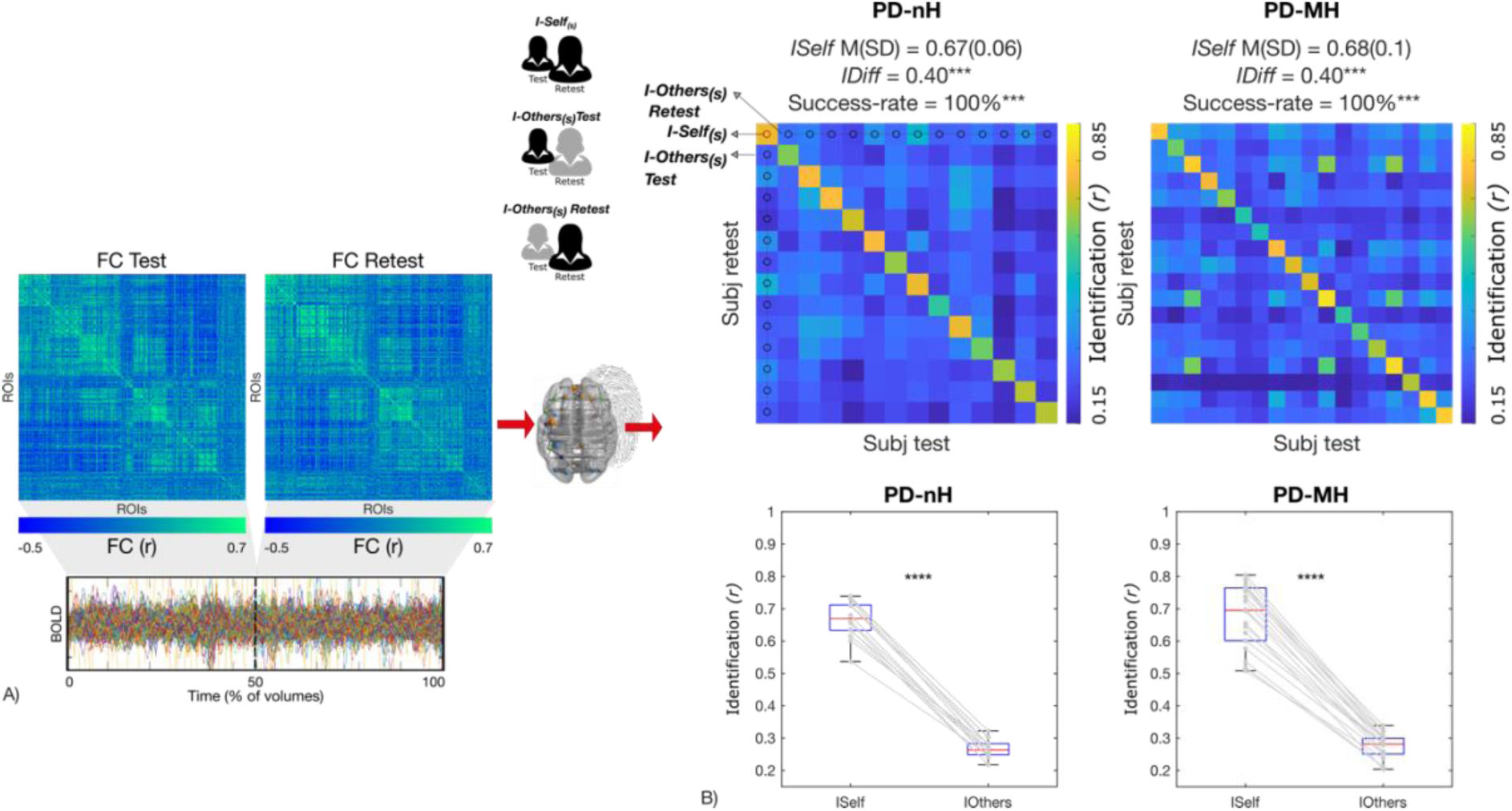
Within and between-subjects variability in functional connectivity. A) Functional connectome estimation at test (first half of the volumes) and re-test (second-half of the volumes). B) Identifiability matrices show within-(*ISelf*) and between-subjects (*IOthers*) test-retest reliability as Pearson correlation coefficient in PD patients without hallucinations (PD-nH) and PD patients with minor hallucinations (PD-MH). Individuals’ *ISelf* and *IOthers* are displayed, respectively, in the diagonal and off-diagonal elements of the matrix. The average ISelf, IDiff and Success-rate were similar in the three groups and IDiff and Success-rate differed at p < .001 (***) from random distributions. Boxplots shows that *ISelf* was significantly higher (paired-sample t-test, p < .0001****) than *IOthers* in all individual cases in both PD-nH and PD-MH.

**Figure 2.**
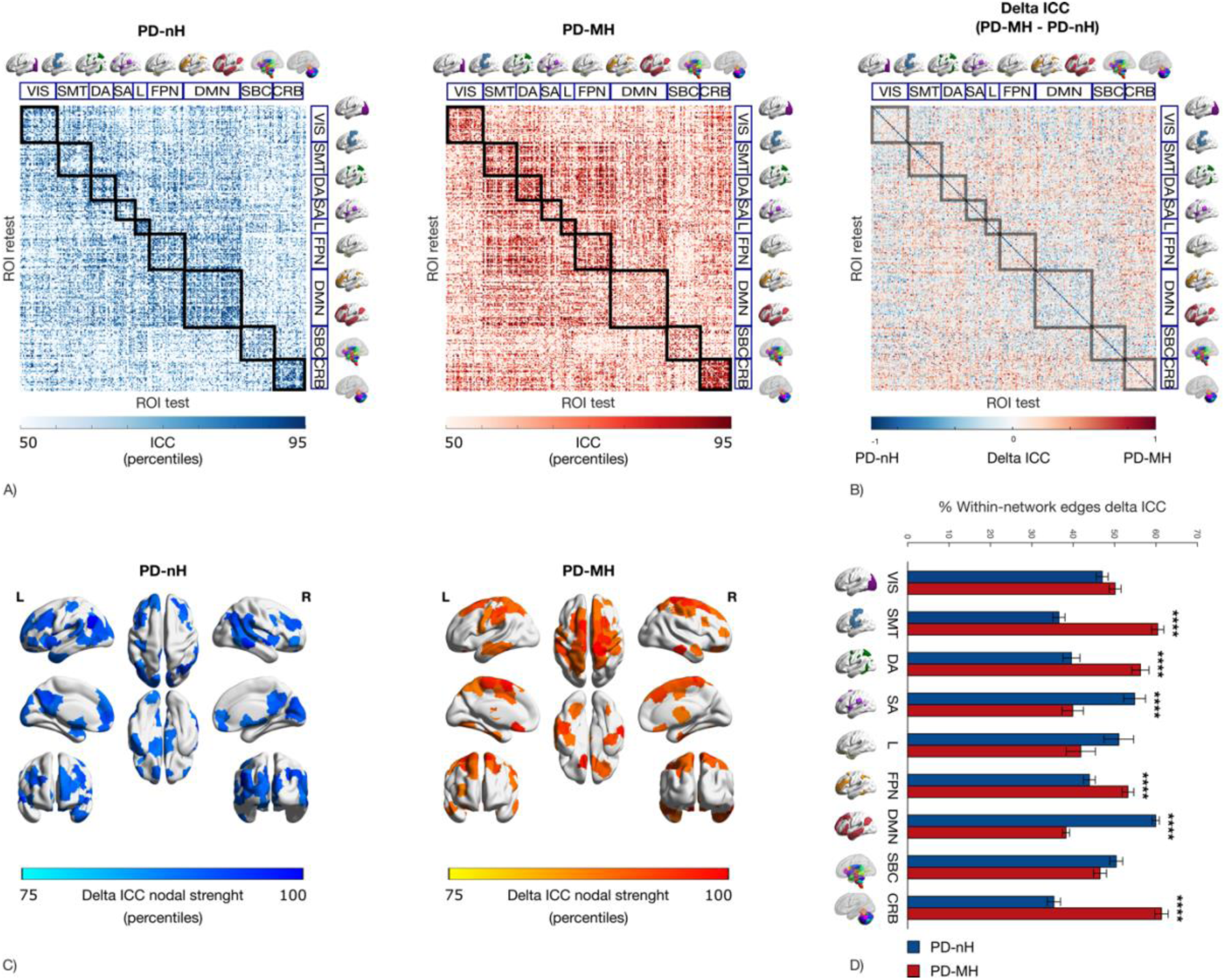
FC patterns making subjects identifiable differ in PD-MH and PD-nH. A) Edgewise intra-class correlations (ICC) matrices for Parkinson’s Disease patients without hallucinations (PD-nH) and with minor hallucinations (PD-MH). ICC quantifies within-subjects similarity between test and retest for each edge (FC between 2 regions). Here we show that a different configuration of the edges with highest ICC in PD-nH vs. PD-MH. We display edges with ICC between the 50^th^ and the 95^th^ percentile, across the 7 resting-state networks (RSN), subcortical and cerebellar regions. B) Delta ICC matrix showing the differential ICC (ICC _PD-MH_ – ICC _PD-nH_) across the 7 RSNs, subcortical and cerebellar regions. VIS=Visual Network; SMT=Somatomotor Network; DA=Dorsal Attention Network; SA=Salience Network; L=Limbic Network; FPN=Fronto-Parietal Network; DMN=Default-Mode Network; SBC=Subcortical regions; CRB=Cerebellum. C) Brain renders show the nodal strength of each group ICC map, masked for delta ICC value for PD-nH and PD-MH group. Nodal strength was computed as average of edge weights for each ROI. We here display only the nodes > 75th percentiles. D) Bar plot show percentage of edges in delta ICC matrix for each of the 7RSNs subcortical and cerebellar regions separately for each group. Comparisons across groups are done using Chi-Square and corrected for multiple comparisons (FDR). **** = p< .0001.

**Figure 3.**
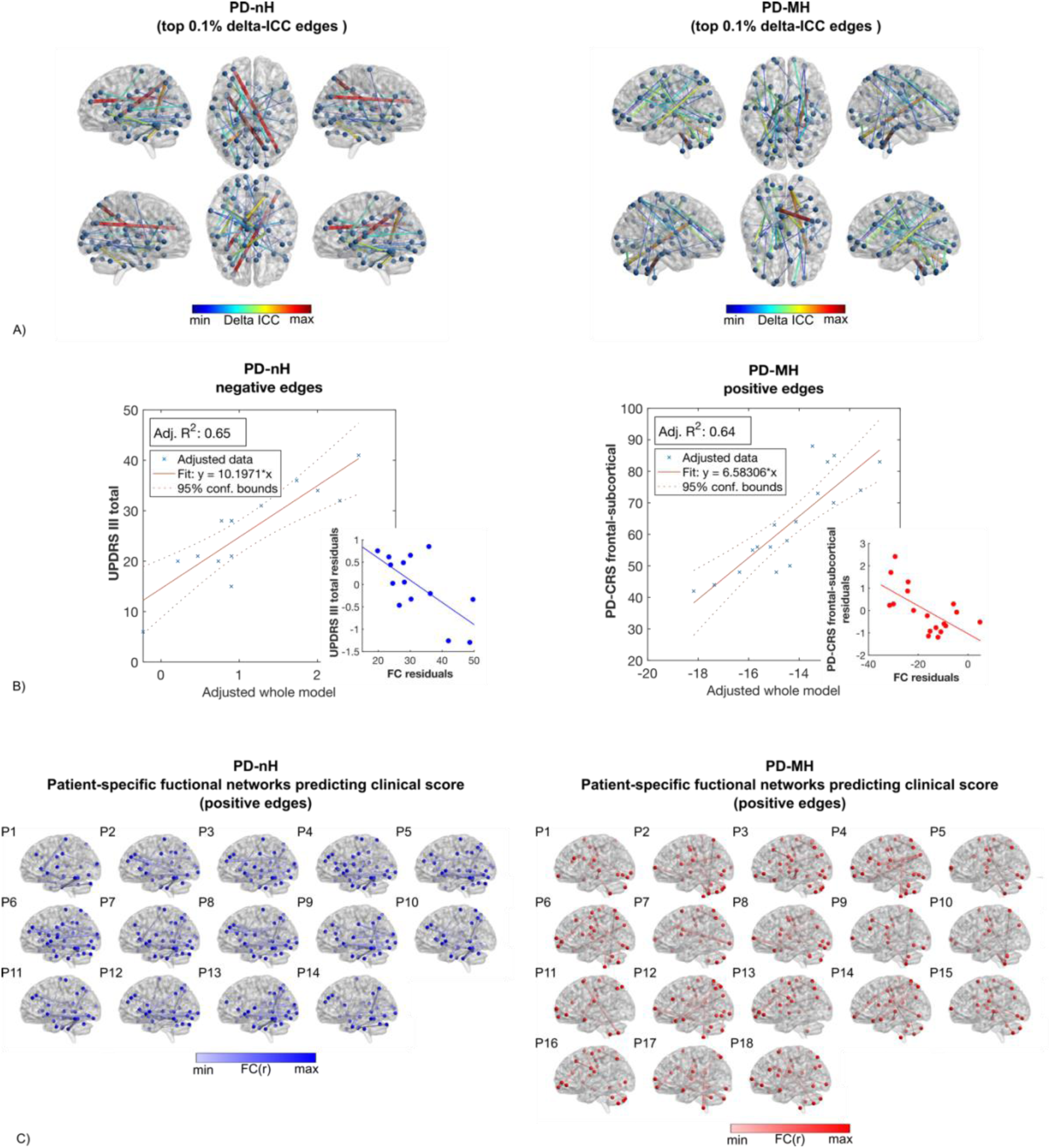
Functional connections with high fingerprint predict clinical scores. A) Brain renders showing the top 0.1% delta ICC edges for each group. The top delta ICC matrices were used as masks to isolate the functional connections that maximise distinguishability in fingerprint between the two groups in each individual functional connectome. C) Large scatterplot: linear regression models where clinical scores were significantly predicted by functional connectivity in the top delta ICC edges (Clinical score ∼ FC _top delta ICC group_ + Age + YoE + Disease duration). Small scatterplot: correlation between residuals of the clinical score and functional connectivity to show directionality of the association. D) Patient’s specific brain renders showing functional connections or edges with high fingerprint predictive of clinical score.

**Figure 4.**
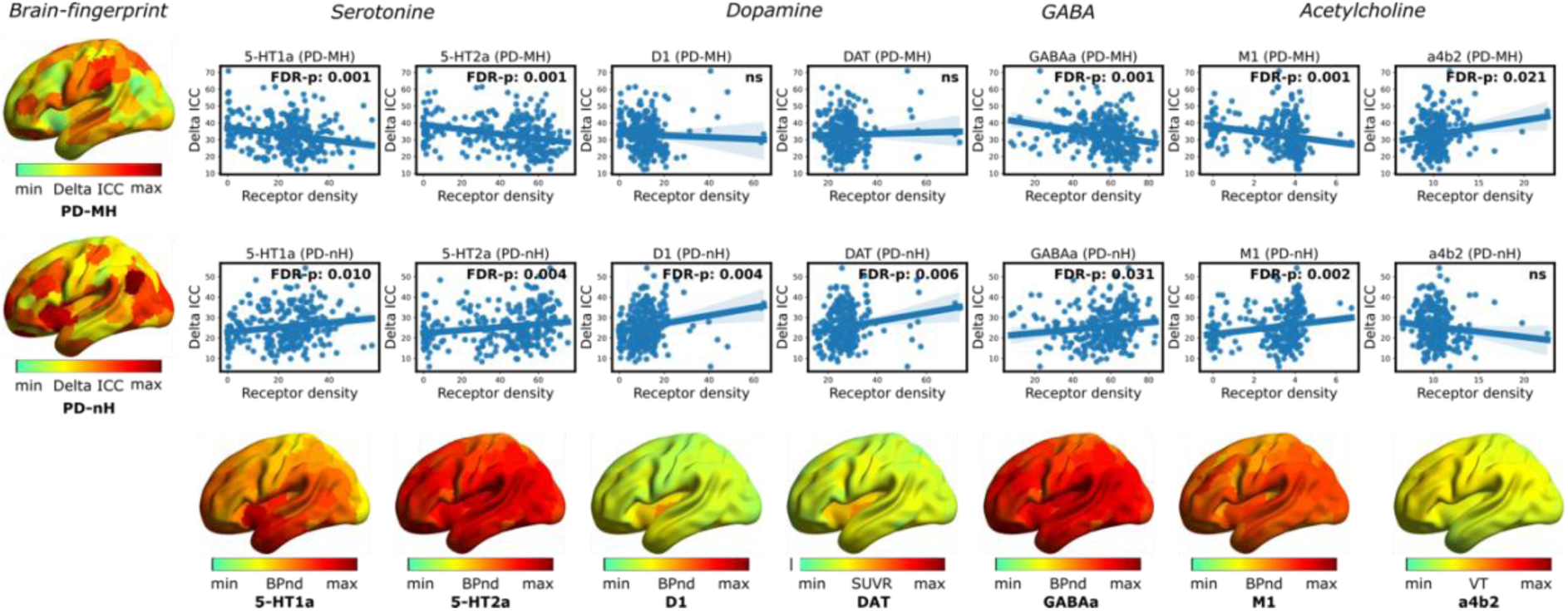
Brain-fingerprint aligns with different neurotransmitter systems in PD-nH and PD-MH. Scatterplots show Pearson’s spatial correlations between the group-wise delta ICC map and neurotransmitter receptor densities for dopamine, serotonin, GABA, and acetylcholine. Reported p-values are FDR-corrected (ns = FDR-p < .05). Serotonin receptors include 5-HT1a and 5-HT2a. Dopamine receptors include D1 and D2 (D2 non-significant in both groups, so not displayed), and DAT refers to the dopamine transporter. GABA receptor refers to GABAa. Acetylcholine receptors include M1 (muscarinic receptor) and α4β2 (alpha-4 beta-2 nicotinic receptor). Abbreviations: BPnd = binding potential; SUVR = standardized uptake value ratio; VT = free-fraction corrected distribution volume.

### Whole-brain within-sessions brain fingerprint

Investigating *whole-brain* fingerprint, we found a 100% *Success-rate* in both groups, demonstrating that each individual can be correctly identified based only on their FC, even within the same group (Fig. 1B, *ISelf* > *IOthers*: p<.001). *IDiff* was equally high (0.40) in PD-MH and PD-nH, indicating strong individual brain discriminability. Test-retest reliability (*ISelf*) was high in both groups, with no significant group differences after controlling for nuisance variables [ANOVA, 5000 permutations; F(1,26)=0.1, p=.766; PD-nH: M(SD)=0.67(0.06); PD-MH: M(SD)=0.68(0.1); Fig. 1B].

Similarly, between-subjects similarity (*IOthers*) showed no group differences [ANOVA, 5000 permutations; F(1,26)=0.95, p=.0.339; PD-nH: M(SD)=0.27(0.03); PD-MH: M(SD)=0.27(0.04)]. See Table 2 for full statistics. Permutation testing confirmed that *IDiff* and *Success-rate* significantly deviated from null distributions (p <.0001) in both PD subtypes (Fig. 1B). In sum, these findings demonstrate that fingerprinting can be applied in PD and that PD patients can be individually identified with high accuracy based solely on within-scan brain connectivity pattern, independent of clinical status, and significantly above surrogate null models.

### Spatial specificity of brain fingerprint

Analysing the local properties of the brain fingerprint using edgewise intra-class correlation (ICC, Amico & Goñi, 2018a), we observed that the functional connections with the highest fingerprint exhibited different spatial distributions in PD-nH and PD-MH (Fig. 2A). To identify ICC edges that distinguished the two groups, we computed an edge-based differential identifiability (ΔICC) matrix (i.e., ICC _PD-MH_ – ICC _PD-nH_; Fig. 2B, see ‘Between-groups differential ICC’). These edges were then mapped onto the cortical surface as nodal strength (Fig. 2C) and computed across each of the 7RSN, subcortical and cerebellar regions (Fig. 2D). This analysis revealed differences between PD-MH and PD-nH: PD-MH showed the highest identifiability in the somatomotor, dorsal attention (DAN), and fronto-parietal (FPN) networks, as well as in cerebellar regions (Fig. 2D). In contrast, PD-nH exhibited higher identifiability in the SAL and DMN (Fig. 2D). In sum, these findings indicate that the brain fingerprint is expressed through distinct functional connections in PD patients with (PD-MH) and without (PD-nH) minor hallucinations.

### Functional connections with high fingerprint predict clinical scores

To identify ICC edges that maximally distinguished the two groups, we selected the top 0.1% edges for each. Their spatial distribution on the cortical surface is shown in Fig. 3A for both groups separately. These top ICC edges were then used to isolate functional connections with the highest edgewise differential identifiability in each individual’s functional connectome, from which we derived a sum score separately for positive and negative connections. Using linear models, we tested whether this functional connectivity score predicted clinical scores of (i) motor symptoms severity (using UPDRS III total score), (ii) frontal-subcortical executive functioning (using PD-CRS frontal subcortical score) and (iii) posterior-cortical functions (using PD-CRS posterior score), separately in each group.

In PD-nH, negative connectivity in high-ICC edges predicted UPDRS III total score (Adjusted R² = 0.65; p = .003; Fig. 3B), indicating that more significant disconnection of fingerprinting hubs (individual brain renders, Fig. 3C) was associated with lower motor impairment. This differed for PD-MH, in whom functional connectivity predicted the PD-CRS Frontal Subcortical neuropsychological score (Adjusted R² = 0.64; p = .010; Fig. 3B), with higher connectivity between fingerprinting hubs (Fig. 3C) linked to less executive impairment. No significant associations were found for the PD-CRS posterior score in neither group, which was consistent with the early disease stage.

### The cortical topography of brain-fingerprint aligns with different neurotransmitter systems in the patients with and without minor hallucinations

We assessed whether the topography of each group’s brain fingerprint aligned with the cortical distribution of dopamine, serotonin, GABA, and acetylcholine receptors, given their role in the neurobiology of hallucinations (Fig. 4). We found that in PD-nH, the fingerprint was positively associated with dopamine receptor density (D1: FDR-p = .006; DAT: FDR-p = .007), while no significant association was found between fingerprint and any dopamine receptor density in PD-MH. In addition, in PD-MH, regions with high fingerprint exhibited lower receptor density across most tested neurotransmitter systems implicated in hallucinations: serotonin (5-HT1A: FDR-p = .001; 5-HT2A: FDR-p = .001), GABA (GABAa: FDR-p = .001), and the muscarinic acetylcholine receptor (M1: FDR-p = .001). Again, PD-nH showed a different pattern, with high-fingerprint regions associated with higher receptor density for serotonin (5-HT1A: FDR-p = .011; 5-HT2A: FDR-p = .032), GABA (GABAa: FDR-p = .032), and the muscarinic acetylcholine receptor (M1: FDR-p = .002). The nicotinic acetylcholine receptor data (α4β2) revealed either no significant association (PD-nH) or higher density in regions with high fingerprint (PD-MH: FDR-p = .021).

## Discussion

In this study, we analysed for the first time the features of brain-fingerprints using resting-state fMRI in patients with Parkinson’s disease (PD) with minor hallucinations (PD-MH) and without any hallucinations (PD-nH). First, we observed that brain-fingerprints are preserved in the presence of PD, irrespective of our patients’ hallucination status (Fig. 1). Second, we identified the spatial features of brain-fingerprints that maximally distinguished the two groups (Fig. 2), which in turn allowed us to identify patient-specific functional networks that predicted clinical traits at the individual level and distinguishing cognitive from motor symptoms in PD-MH vs. PD-nH, respectively (Fig. 3). Finally, the spatial features of the brain-fingerprints were associated with the cortical density of neurotransmitter systems linked to hallucinations, again with distinct patterns in patients with and without MH.

Our results demonstrate that individual functional connectivity patterns contain stable, subject-specific features, enabling patient identification in PD. These findings extend previous evidence from MEG studies in PD ^46,47^ and align with fMRI studies in other neurodegenerative disorders ^45^. The persistence of brain fingerprinting in neurodegenerative conditions is a non-trivial finding because it shows that even within a homogeneous group – patients with the same diagnosis, same range of biological alterations and symptoms, and, in this case, belonging to the same subtype with potentially similar clinical trajectories – each individual remains highly distinct. Features embedded in these functional connectomes hold, therefore, great promise for advancing personalized prognostics and treatments by providing objective, data-driven biomarkers to identify individual differences, track disease progression, and may predict variability in therapeutic responses ^49,50^.

In healthy subjects, the most prominent networks for subject identification are the DMN, FPN and SAL ^36,37^. In the present study, PD patients without hallucinations –who are less likely to experience cognitive decline ^12^ – maintain high identifiability relying upon the DMN, FPN and SAL (Fig. 2A). This differs for PD-MH patients, who have been considered to have a more severe and rapidly advancing form of PD, often associated with cognitive impairments ^11^. Here, we report for PD-MH patients a significant reduction in DMN and SAL identifiability (Fig. 2B and 2D), suggesting a more substantial loss of subject-specific features in these two key networks, potentially reflecting an early fragility/impairment of this group. Of note, over-reliance on the DMN and SAL network, as only observed in the present PD-MH group, has been implicated in hallucinations, but only for complex visual hallucinations ^24–26^ that arise much later in the disease course ^11^. Our findings reveal a similar distinction using a different methodological approach, but at an earlier disease stage and using a personalized patient-specific framework.

Concerning the FPN, we found that PD-MH patients showed a differential FPN fingerprint, with more connections exhibiting higher a fingerprint in PD-MH vs. PD-nH, although there was a preserved high FPN fingerprint in both groups (Fig. 2B). The preserved high fingerprint in the FPN in both groups (Fig. 2B) may reflect that executive functioning is still largely intact in this sample of patients (tested early in disease course), whereas the increased FPN fingerprint only in PD-MH is compatible with a more cortical-diffuse or advanced form of PD, as argued elsewhere ^9,11^. We note that the present patient groups did not suffer from significant executive impairment ^51,52^ and had normal PD-CRS frontal-subcortical scores that did not differ [63.3(14.9) for PD-MH and 64.4(14.8) for PD-nH, cf. Table 1].

**Table 1.** Demographics and clinical variables.

|  | PD-MH | PD-nH | p value | p value sig. |
| --- | --- | --- | --- | --- |
| N | 18 | 14 | - |  |
| Sex (%) | 44.4 | 28.6 | 0.471 | ns |
| Age M(SD) | 70.4(5.5) | 65.8(7.8) | 0.07 | ns |
| YoE M(SD) | 12.5(4.6) | 11.6(4.9) | 0.618 | ns |
| Motor symptoms severity: UPDRS III - M(SD) | 21.9(8.6) | 25.8(9.2) | 0.238 | ns |
| Cognitive functions: PD-CRS (Frontal max=104) | 63.3(14.9) | 64.4(14.8) | 0.848 | ns |
| Cognitive functions: PD-CRS (Posterior max=30) | 28.6(1.4) | 28.6(1.7) | 0.875 | ns |
| Disease duration: years - M(SD) | 5.2(3.8) | 4(2) | 0.253 | ns |
| Daily dose of L-dopa | 697.2(311.4) | 601.1(311.9) | 0.394 | ns |
| Dopamine agonist dose | 143.7(117.9) | 153.6(139.9) | 0.833 | ns |

*Legend:* PD-MH=Parkinson's disease with minor hallucinations; PD-nH= Parkinson's disease with no hallucinations. M=Mean; SD=Standard deviation; YoE=years of education; UPDRS=Unified Parkinson's disease rating scale; PD-CRS=Parkinson's disease - cognitive rating scale; p-value was estimated using Fisher's test for sex, and two-samples t-test or its non-parametric equivalent, i.e., Wilcoxon signed-rank test for the remaining variables. ns=not significant, $p>0.05$ ; \* $p\leq0.05$ ; \*\* $p\leq0.01$ ; \*\*\* $p\leq0.001$ ; \*\*\*\* $p\leq0.0001$ .

The somatomotor network is typically not among those with the highest fingerprint in healthy subjects; however, in PD-MH, it stands out as one of the networks with the highest ICC, both within-network and in its connections with the DAN, SAL, limbic, and FPN networks (Fig. 2B). This underscores the importance of the somatosensory and motor functions in the pathogenesis of MH, including presence hallucinations, which have recently been linked to specific somatomotor processes in PD patients ^33^ and in healthy subjects in whom presence hallucinations have been induced experimentally using a somatomotor conflict (including specific tactile, motor, proprioceptive signals)^53^, associated with activation of primary motor and premotor cortex (cf. Figure 2E in ^33^).

These differential results across the two patient groups were further supported by the finding that these differences allowed the identification of patient-specific functional networks predicting different clinical traits in the two groups. In PD-MH patients, stronger connectivity in these regions was associated with lower scores for frontal-subcortical functions (PDCRS score). Thus, although early PD-MH patients (including the present sample) generally have normal cognitive scores, they may show subtle sub-clinical deficits in frontal-subcortical functions (including executive and attentional functions ^9,54^). Standard resting-state fMRI analyses at the group level reported that altered connectivity in a priori defined fronto-parietal connections in PD-MH was associated with frontal-subcortical scores^33^. The present study significantly extends these findings by showing that individualized functional connectivity patterns—rather than group-level effects— can be derived in an entirely data-driven manner and are detected in PD-MH patients with early sub-clinical cognitive alterations. This concerned only frontal-subcortical functions and was not observed in PD-nH patients. Taken together, these results suggest that MH in PD involves a dysfunction of frontal-executive and somatomotor networks, distinguishing PD patients with versus without MH. In contrast, for PD-nH patients – who typically show no or less rapid cognitive decline – we found that greater disconnection in regions with high fingerprint was associated instead with lower motor functions (UPDRS-III total score), further distinguishing a motor-only form of PD from a motor-cognitive form of PD. This may allow early distinction between both groups and different therapeutical strategies, as recent work has revealed different clinical trajectories and different sensitivities to disease-modifying treatments ^3^.

We argue that these findings are particularly noteworthy as they could open new avenues for individualized neuromodulation targets. While the current symptom-specific approach – where stimulation is personalized based on the most burdensome symptoms using group-derived targets – has significantly advanced the field ^55^, integrating an individualized network-based perspective could further refine treatment. Rather than targeting symptoms alone, this approach would tailor stimulation to the specific brain networks associated with each symptom in a particular patient. If these findings hold in other cohorts, this could pave the way for defining an algorithm for brain-network personalized treatment.

Finally, the present receptor data show for PD-nH a positive association with D1 and DAT, suggesting that D1 dopaminergic activity plays an important role in this group of non-hallucinating patients. Confirming clinical and brain fingerprinting differences between the groups, no such association was found for any dopaminergic receptor (D1, D2, DAT) and brain fingerprint in PD-MH. While the absence of a correlation between the regional density of dopamine receptors and brain fingerprint in PD-MH does not rule out an indirect dopaminergic influence on minor hallucinations, our findings emphasize the role of other non-dopaminergic neurotransmitters in the development of PD-related hallucinations, similarly to ^56^. Namely, PD-MH patients exhibited lower serotonin (5-HT1A, 5-HT2A), GABAa, and muscarinic M1 receptor densities in high-fingerprint regions, while PD-nH showed the opposite pattern, with higher densities in these areas. Since the neurotransmitter maps are derived from healthy subjects, interpreting the directionality of these associations is complex. Previous studies have found increased 5-HT2A receptor binding in post-mortem PD patients with hallucinations in the inferolateral temporal cortex ^57^, and in vivo studies have shown higher receptor binding in the visual, dorsolateral, medial-orbitofrontal cortices, and insula in PD patients with hallucinations ^58^. Similarly, other research has found reduced occipital GABAa in PD patients with visual hallucinations ^59^, and other evidence has shown that blockade of muscarinic receptors can induce hallucinations ^60^ and grey matter volume reduction in cholinergic areas ^61^. These regions do not overlap with the high-fingerprint areas where we observed lower receptor densities in PD-MH patients. Although speculative, this suggests that high-fingerprint regions may be relatively spared from disease-related alterations and that the temporal connectivity stability that is captured by brain-fingerprint ICC could represent a compensatory mechanism.

The present study has some limitations. First, while the sample size is relatively limited, this study stands out as a rare opportunity to investigate a cohort stratified for minor hallucinations. Such databases are scarce, making this work a valuable step toward understanding brain-fingerprint in this specific group of patients and early in the disease course. Second, this study divided the same scanning session into two in order to estimate brain identifiability, addressing challenges in obtaining two separate test-retest sessions in clinical cohorts as in ^45^. While this approach been shown to yield similar results to between-session data ^37^, future research will need to replicate our findings. Finally, while our results provide valuable insights into the features of brain-fingerprint in PD patients with minor hallucinations, it would be particularly interesting to examine how these features evolve over time, if they can predict later clinical cognitive impairments, and whether the observed fingerprinting patterns change as new symptoms appear.

In conclusion, our study provides novel evidence that fMRI brain fingerprints are preserved in PD and can distinguish between patients with and without MH. These differential fingerprinting features linked to distinct neurobiological substrates and allowed the identification of patient-specific functional networks associated with clinical traits. Notably, we identified patient-specific functional networks associated with sub-clinical cognitive impairment in early-stage PD patients with MH – a subtype associated with cognitive decline and dementia. This fully data-driven framework could, therefore, open avenues for personalized early intervention, prognostic, and therapeutic strategies. Our findings also align with previous research linking MH to somatomotor and fronto-parietal networks, reinforcing the relevance of these cortical networks in PD patients with hallucinations ^9,33^. Future work should explore how these patterns evolve alongside disease progression and whether they can be leveraged as early biomarkers for cognitive decline susceptibility in PD.

## Materials and Methods

Clinical and imaging data were the same used in ^35^.

### Participants

N=32 patients with PD who previously participated in ^35^ where included in the analyses. All patients fulfilled the MDS new criteria for PD ^62^ with minor hallucinations (PD-MH, N=18) and without hallucinations (PD-nH, N=14). MH included presence (N=11), passage (N=10) hallucinations, visual illusions or pareidolias (N=6) and/or a combination of two or more types minor hallucinations (N=9). Demographics are provided in the ‘Statistics and Reproducibility’ section). Further details about inclusion/exclusion criteria and clinical evaluation can be found in ^35^.

### Image acquisition parameters

MRI scans were acquired with a 3T Philips Achieva. T1 weighted scans were obtained using a MPRAGE sequence (TR = 500 ms, TE = 50 ms, flip angle = 8, field of view [FOV] = 23 cm with in-plane resolution of 256 × 256 and 1mm slice thickness). Resting-state functional MRI images were collected using an 8-minute sequence (TR = 2000 ms, TE = 30 ms, flip angle = 78, FOV = 240 mm, slice thickness = 3 mm, 180 volumes).

### Image processing

fMRI data were preprocessed using in-house MATLAB code based on state-of-the-art fMRI processing guidelines ^63–65^, using the open-source MATLAB toobox Apéro (https://github.com/juancarlosfarah/apero). Below follows a brief description of these steps. Structural images were first denoised to improve the signal-to-noise ratio ^66^, bias-field corrected, and then segmented (FSL FAST) to extract white matter, grey matter and cerebrospinal fluid (CSF) tissue masks. These masks were warped in each individual subject’s functional space by means of subsequent linear and non-linear registrations (FSL flirt 6dof, FSL flirt 12dof and FSL fnirt). The following steps were then applied on the fMRI data: BOLD volume unwarping with applytopup, slice timing correction (slicetimer), realignment (mcflirt), normalisation to mode 1000, demeaning and linear detrending (MATLAB detrend), regression (MATLAB regress) of 18 signals: 3 translations, 3 rotations, and 3 tissue-based regressors (mean signal of whole-brain, white matter (WM) and cerebrospinal fluid (CSF), as well as 9 corresponding derivatives (backwards difference; MATLAB). We tagged high head-motion volumes on the basis of three metrics: frame displacement (FD, in mm), standardised DVARS ^67^ (D referring to temporal derivative of BOLD time courses, VARS referring to root mean square variance over voxels) as proposed in ^64^, and SD (standard deviation of the BOLD signal within brain voxels at every time-point). The FD, DVARS were obtained with fsl_motion_outliers and SD vectors with MATLAB. Volumes were motion-tagged when FD > 0.3 mm and standardised DVARS > 1.7 and SD > 75th percentile +1.5 of the interquartile range, as per FSL recommendation ^68^. All subjects had ≤30% motion-tagged volumes. There was no significant difference across groups in the percentage of tagged volumes [p=.289], nor in the average FD [p=.593] or DVARS [p=.418]. There was no difference across test and retest in the percentage of tagged volumes [p=.661] nor in the average FD [p=.081] nor DVARS [0.899].

A bandpass first-order Butterworth filter [0.01 Hz, 0.15 Hz] was applied to all BOLD time-series at the voxel level (MATLAB butter and filtfilt). The choice of the bandpass filter was aligned with previous works ^69^ where the choice of the filtering proved to be meaningful to capture the effect of brain fingerprints in the temporal domain, and in the analogy between MEG and fMRI fingerprints^70^, and with respect to the relationship between brain fingerprints and structure-function coupling ^71^.

The first three principal components of the BOLD signal in the WM and CSF tissue were regressed out of the grey matter (GM) signal (MATLAB, pca and regress) at the voxel level. A whole-brain data-driven functional parcellation based on 278 regions including cortical and subcortical areas as obtained by ^72^, was projected into each subject’s T1 space (FSL flirt 6dof, FSL flirt 12dof and finally FSL fnirt) and then into the native EPI space of each subject. We also applied FSL boundary-based-registration ^73^ to improve the registration of the structural masks and the parcellation to the functional volumes.

### Functional Connectivity and whole-brain within-session brain-fingerprint

We estimated individual FC matrices using Pearson’s correlation coefficient between the averaged signals of all region pairs. The resulting individual FC matrices were composed of 278 cortical nodes, as obtained by ^72^. Finally, the resulting functional connectomes were ordered according to seven cortical resting state networks (RSNs) as proposed by ^74^, plus subcortical and cerebellar regions (similarly to ^75^, see also Fig. 1A).

We estimated *within-session* identifiability or fingerprinting, following the approach proposed. This method involves splitting the fMRI times series in halves and enables quantification of *within-session connectome fingerprints.* Previous work has demonstrated that this method produces very similar results to those obtained from data acquired across separate sessions (i.e., between-sessions fingerprint) in healthy subjects from the Human Connectome Project (HPC) (Fig. S3 in ^48^). Although *within* and *between-sessions fingerprinting* held similar results, they are different approaches to quantifying the brain-fingerprint, and this should be compared in future studies. However, we note that there are currently no clinical datasets of PD patients with minor hallucinations available that include two fMRI sessions acquired within a short-time gap (i.e., within around one or two weeks). Therefore, *within-session fingerprint* is currently the only method available for estimating brain-fingerprint in PD patients with minor hallucinations. In this current study, we estimated identifiability across the first half (test) and second half volumes (retest) within the same scan. Recent work has shown that a good level of identifiability across the different resting state networks can be reached from around 200s (see Fig 4B, in ^69^). In this work, each test and retest session had 90 volumes with a TR of 2s, therefore providing sufficient data for achieving a good success rate and identifiability across the entire brain.

At the whole-brain level, the fingerprint was calculated for each subject 𝑠 as test-retest similarity between FCs (cf. Fig 1B; we called this metric *ISelf*).

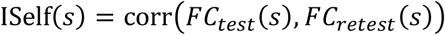

Then, for each subject 𝑠 we computed an index of the FCs similarity with the other subjects 𝑖 in their group (*IOthers*), where 𝑁 is the total number of subjects in each group:

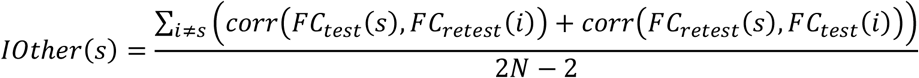

A second metric, *IDiff* (Fig. 1B), provides a group-level estimate of the within-(*ISelf*) and between-subjects (*IOthers*) test-retest reliability distance, where 𝑆𝑢𝑏𝑗 is the set of subjects:

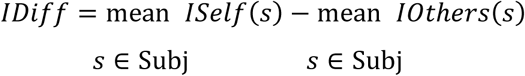

Finally, we measured the Success-rate ^36^ of the identification procedure as the percentage of cases with higher within- (*ISelf*) vs. between-subjects (*IOthers*) test-retest reliability. These metrics have been introduced and estimated in patients ^76^ and healthy populations ^48^ in previous work.

### Spatial specificity of brain fingerprint: edge-wise intra-class correlation

Spatial specificity of FC fingerprints was derived using edgewise intra-class correlation (ICC) with one-way random effect model according to ^77^ (cf. Fig. 2A):

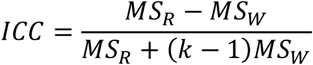

Where 𝑀𝑆_𝑅_= mean square for rows (between the subjects); 𝑀𝑆_𝑊_ = mean square for residual sources of variance; 𝑘 = sessions. ICC coefficients quantify the degree of similarity between observations/measures and find high applicability in reliability studies ^78^. The higher the ICC coefficient, the stronger the agreement between two observations. Here we used this metric, as in previous work ^48,69^, to quantify the similarity between test and retest for each edge (FC between 2 regions). A high ICC indicates that a larger proportion of the variance across test and retest is due to differences between the subjects, rather than differences between test and retest or random error. A low

ICC, on the other hand, indicates that there is more variability due to differences between test and retest or random error, than due to differences between subjects. In other words, the higher the ICC of an edge, the more that edge connectivity is similar for each subject across test and retest, as well as the variability across subjects, i.e., the higher the ‘fingerprint’ of that edge.

The ICC has been commonly used in the functional connectivity literature to assess reliability ^48,79–82^. A few strengths of the ICC include: 1) the ability to assess absolute agreement in repeated measurements of an object (unlike, for example, Pearson’s correlation, wherein variables are scaled and cantered separately), 2) the ability to explicitly model multiple known sources of variability (e.g., scanner, brain response, head motion), and 3) comparing within and between variability across the objects of measurement. Depending on whether and how sources of error (or “facets”; e.g., scanner) may be specified, one of three ICC forms may be used ^83^. In brief, usage is as follows: ICC(1,1) is used to estimate agreement in exact values when sources of error are unspecified; ICC(2,1) is often referred to as “absolute agreement” and is used to estimate agreement in exact values when sources of error are known (e.g., repeated runs) and modelled as random; and ICC(3,1) is often referred to as or “consistent agreement” and is used to estimate agreement in rankings when sources of error are known and modelled as fixed (resulting in a mixed effects ANOVA). In this paper we used ICC(1,1), following previous works^48^, because variability in fMRI data can come from different known (e.g., scanner, head motion) and unknown sources. In the statistics literature the ICC is akin to a measure of discriminability and is commonly categorized as follows: poor <0.4, fair 0.4–0.59, good 0.6–0.74, excellent ≥0.75^84^. In this paper, this categorization was taken as reference for the thresholds on the ICC matrices, which was set at 0.6 (good^84^).

Edge-wise ICC was computed for all possible edges and for each group separately, with the aim to quantify the edges-wise functional connectivity fingerprint, distinctive of each clinical group. In order to control for sample size differences across groups, bootstrapping was used to accurately estimate edgewise fingerprints: for each group, ICC was calculated across test and retest for subsets of randomly chosen N=10 subjects, across 1000 bootstrap runs, and then averaged within each group (Fig. 2A). Bootstrapping was performed in MATLAB using an in-house function.

### Between-groups differential ICC

In these analyses, we aimed to identify edges that were group-specific, meaning those that contributed to differences in fingerprint between the two groups. To achieve this, we first computed a delta ICC matrix to capture differential identifiability, defined as:

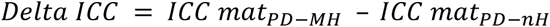

As a result, the delta ICC matrix (Fig. 2B) contained positive values for functional connections with higher ICC in PD-MH and negative values for those with higher ICC in PD-nH. Values close to zero indicated similar ICC across both groups, suggesting individual variability that did not contribute to distinguishing features between them. We identified group-specific ICC edges by selecting the positive or negative delta-ICC edges, for PD-MH and PD-nH respectively. We then computed nodal strength of these group-specific ICC matrices to visualise ICC hubs on the cortical surface (using BrainNet ^85^). To highlight most prominent hubs, we applied a threshold, displaying only brain regions with nodal strength above the 75th percentile (Fig. 2C). Finally, we examined the distribution of the group-specific ICC edges across functional networks. Specifically, we computed the percentage of edges per network separately for PD-MH and PD-nH and tested for group differences using a Chi-Square test, applying FDR correction for multiple comparisons (Fig. 2D).

### Predicting clinical scores from brain fingerprint

In these analyses, we tested whether clinical scores could be predicted from individuals’ functional connections that most strongly distinguished the two groups in terms of fingerprint. To identify these key connections, we selected in the top 0.1th percentile edges for each group-specific ICC matrix. These edges were visualized on the cortical surface using BrainNet (Xia et al., 2013) (Fig. 3A). The binarized matrices served as masks to extract high-fingerprint edges from each individual’s functional connectome (Fig. 3C). For each individual, we separated positive and negative functional connections to prevent mathematical cancellation of connectivity values. We then computed two summed connectivity scores: one for positive and one for negative connections. Next, we fitted linear regression models (using MATLAB’s fitlm function) to predict clinical scores based on individual functional connectivity (Fig. 3B). Each model included a clinical score as the dependent variable and a functional connectivity value for each individual as the predictor, while controlling for age, years of education, and disease duration.

The clinical scores analysed included the Unified Parkinson’s Disease Rating Scale (UPDRS) III total, a global measure of motor symptom severity ^86^. We also examined the PD-CRS Frontal Subcortical score, a composite measure of executive function that includes sustained attention, working memory, verbal fluency, clock drawing, and verbal memory tasks. This score is sensitive to mild cognitive impairment in PD ^87^ and has been linked to neural alterations specific to PD patients with MH ^9,33^. In contrast, the PD-CRS Posterior score, which assesses confrontation naming and clock copying, is reportedly altered later, in the transition to dementia ^87^.

#### Association between cortical topography of brain-fingerprint and neurotransmitter systems

In these analyses, we looked at which neurotransmitter systems showed topographical correspondence with the brain fingerprint topography (Fig. 4). For each group, we estimated the Pearson’s spatial correlation between the group-wise delta ICC map derived from nodal strength of the group-specific ICC matrix and each of the normative atlas maps of the neurotransmitters’ receptors of dopamine, serotonin, GABA and acetylcholine given their implication in the neurobiology of hallucinations. Normative neurotransmitter density data are available from neuromaps (https://github.com/netneurolab/neuromaps) ^88^.

For serotonin, 5-HT1A and 5-HT2A receptor densities were derived from PET tracer binding potential (BPnd) in 95 healthy subjects ^89^. For dopamine, D1 and D2 densities were derived from PET BPnd in 13 ^90^ and 7 ^91^ healthy subjects, respectively, with D2 measured across two retest sessions. Dopamine transporter (DAT) density was based on SPECT standardized uptake value ratio (SUVR) in healthy volunteers ^92^. GABAa density was derived from PET BPnd ^92^. For acetylcholine, muscarinic M1 receptor density was obtained from PET BPnd in 6 healthy subjects across test-retest sessions ^93^, and nicotinic α4β2 receptor density was derived from PET free-fraction corrected distribution volume (VT) in 8 healthy subjects ^94^.

Each receptor map was parcellated using the 278 parcellation from ^72^ to match the ICC maps. We used non-parametric permutation test (permtest_metric from neuromaps ^88^) to assess the Pearson’s correlation between the receptor map and the ICC map, generating a null distribution via 10,000 permutations. A two-tailed p-value was computed as the proportion of permuted correlations equal to or more extreme than the observed correlation. To correct for multiple comparisons, we applied the false discovery rate (FDR) correction using the Benjamini-Hochberg procedure (multipletests function from the python statsmodels package).

### Statistics and reproducibility

The statistical tests reported in this manuscript are two-sided and performed in RStudio 2022.07.2 ^95^, Python 3.9.18 and MATLAB R2022a ^96^. Normality assumptions were checked prior the analyses using Shapiro-Wilk Normality Test (Shapiro_test R function, rstatix package). When normality was not met, non-parametric equivalent was used, as detailed below.

## Demographics

N = 32 subjects with PD were included, of which N=18 with minor hallucinations (PD-MH) and N=14 without hallucinations (PD-nH). Differences across groups were tested using Fisher’s test for sex, and two-samples t-test or its non-parametric equivalent, i.e., Wilcoxon signed-rank test for the remaining variables. There were no differences across groups for sex (p=.471), age (p=.070), years of education (p=.618), motor symptoms severity (UPDRS III total: p=.238), cognitive functions (PD-CRS Frontal Subcortical: p=.848; PD-CRS Posterior: p=.875), disease duration (p=.253) and daily dose of L-dopa (p=.394) and dopamine agonist dose (p=.833; cf. Table 1).

### ISelf and IOthers

*ISelf* and *IOthers* were normally distributed in each group. Analyses were performed using MATLAB and RStudio. To compare *ISelf* vs. *IOthers* in each group we performed paired-sample t-test (ttest, MATLAB 2022a). Then, we used one-way ANOVAs to test the effect of group on *ISelf* and *IOthers* separately after checking for nuisance variables, with 5000 permutations to control for sample size differences (using aovperm from permuco R-package)^97^ (Table 2). For *ISelf*, the nuisance variables were age, sex, years of education (YoE), and the difference in motion between the test and retest scans, as absolute difference between FD (*ISelf* ∼ Group + Age + Sex + YoE + delta FD). For *IOthers*, the nuisance variables were also age, sex, and YoE, but motion (FD) was considered across the entire acquisition, as *IOthers* is a composite measure across test and retest *(IOthers ∼ Group + Age + Sex + YoE + FD).* Finally, we did a permutation testing analysis to compare *Success-rate* and *IDiff* from 1000 surrogate datasets of random ID matrices against the real value ^70^. Permutation analyses were implemented in MATLAB using in-house function where a permuted version of the ID matrix was built by randomly shuffling its elements.

**Table 2.** Difference across groups in *ISelf* and *IOthers*.

| *ISelf* ~ *Group* + *Age* + *Sex* + *YoE* + $\Delta$ *FD* | | | | | | |
| --- | --- | --- | --- | --- | --- | --- |
| *Anova Table* | SS | df | F | parametric<br>P(>F) | resampled<br>P(>F) | p resampled<br>signif. |
| Group | 0.001 | 1 | 0.1 | 0.768 | 0.774 | ns |
| Age | 0.002 | 1 | 0.3 | 0.573 | 0.570 | ns |
| Sex | 0.008 | 1 | 1.1 | 0.303 | 0.304 | ns |
| YoE | 0.000 | 1 | 0.1 | 0.819 | 0.813 | ns |
| $\Delta$ FD | 0.001 | 1 | 0.2 | 0.661 | 0.666 | ns |

| *IOthers* ~ *Group* + *Age* + *Sex* + *YoE* + *FD* |  |  |  |  |  |  |
| --- | --- | --- | --- | --- | --- | --- |
| *Anova Table* | SS | df | F | parametric<br>P(>F) | resampled<br>P(>F) | p resampled<br>signif |
| Group | 0.001 | 1 | 1.0 | 0.339 | 0.332 | ns |
| Age | 0.002 | 1 | 2.0 | 0.174 | 0.177 | ns |
| Sex | 0.000 | 1 | 0.3 | 0.576 | 0.574 | ns |
| YoE | 0.001 | 1 | 0.4 | 0.517 | 0.515 | ns |
| FD | 0.002 | 1 | 1.3 | 0.257 | 0.256 | ns |

*Legend:* YoE=years of education; $\Delta$ FD=differential frame-wise displacement at test vs. retest; FD=average frame-wise displacement. p-value was estimated using ANOVA with 5000 permutations to control for sample size differences, after controlling for nuisance variables (see Table title for specs about the model). Resampling test using Freedman-Lane to handle nuisance variables and 5000 permutations; ns=not significant > 0.05.

## Inclusion & ethics statement

All subjects gave their informed consent for inclusion before they participated in the study, in accordance with the guidelines of the Declaration of Helsinki. All ethical regulations relevant to human research participants were followed. Patients were recruited at the Movement Disorders Clinic at Hospital de la Santa Creu i Sant Pau (Barcelona) and the study was approved by the local Ethics Committee.

## Data availability

The derived FC matrices and behavioural data necessary to reproduce the main analyses of this study will be made available upon publication in S.S.’s GitHub repository (https://github.com/ss1913/fingerprints_parkinson_mh).

## Code availability

The full code necessary to reproduce the main results and figures will be made available upon publication in S.S.’s GitHub repository (https://github.com/ss1913/fingerprints_parkinson_mh).

## Acknowledgments

This research was supported by the Swiss National Science Foundation [grant n. 188798], CARIGEST SA (Fondazione Teofilo Rossi di Montelera e di Premuda and a second one wishing to remain anonymous) and Parkinson Suisse to O.B.; the EPFL Neuro <u>X</u> Post-doctoral fellowship program to S.S.; the Swiss National Science Foundation [grant n. 221182] and the Leenaards Foundation to F.B.; the Synapsis Foundation to O.B and F.B.; the CIBERNED (Carlos III Institute) and FIS [grant n. PI18/01717] and the Institutode Salud Carlos III (ISCIII), Spain, to J.K.; the PERIS [expedient number SLT008/18/00088] Generalitat de Catalunya to J. P..

## Author contributions

Authors’ contributions according to the CRediT taxonomy (see http://credit.niso.org/ for more information).

*Conceptualization*: S.S. and O.B. (lead); F.B. provided ideas and contributed to the evolution of the overarching research goals and aims.

*Validation*: S.S. *Formal anal*ysis: S.S. *Investigation*: J.P., J.K.

*Resources*: S.S., J.P., J.K., O.B.

*Data curation*: S.S., F.B., J.P., J.K.

*Writing – Original Draft*: S.S. and O.B.

*Writing – Review & Editing*: S.S., F.B., D.V.D.V., E.A., J.P., J.K., O.B.

*Visualization*: S.S.

*Supervision*: O.B.

*Project administration*: S.S., F.B., O.B.

*Funding acquisition*: S.S., O.B. and F.B..

